# Inhibition of miR-22-3p reduces kidney disease associated with systemic lupus erythematosus

**DOI:** 10.1101/512848

**Authors:** Danielle L. Michell, Ashley Faust, Jared L. Moore, Brenna D. Appleton, Michelle Ormseth, Marisol Ramirez-Solano, Quanhu Sheng, Joseph F. Solus, C. Michael Stein, Kasey C. Vickers, Amy S. Major

**Affiliations:** Department of Medicine, Vanderbilt University Medical Center; Department of Pathology, Microbiology and Immunology, Vanderbilt University; Tennessee Valley Health System, Department of Veterans Affairs; Department of Biostatistics, Vanderbilt University Medical Center

**Keywords:** microRNAs, kidney disease, systemic lupus erythematosus, lupus nephritis

## Abstract

Cellular microRNAs (miRNA) have proven to be critical regulators of inflammatory gene expression across many pathways within autoimmunity. Circulating miRNAs serve as a new class of disease biomarkers. Nevertheless, the functional roles of miRNAs, particularly extracellular miRNAs, in systemic lupus erythematosus (SLE) remain poorly understood. Therefore, we aimed to link changes in extracellular miRNAs to lymphocyte gene regulation and the pathophysiology of SLE. Here, we demonstrate that circulating miR-22-3p levels are associated with SLE, and miR-22-3p regulates T and B cell function and SLE-associated kidney disease. Based on high-throughput small RNA sequencing and real-time PCR, extracellular miR-22-3p levels were found to be significantly increased in whole plasma in human SLE subjects. To determine the functional impact of miR-22-3p in SLE, miR-22-3p loss-of-function studies were performed in a mouse model of SLE (B6.*SLE1.2.3)*. We found that *in vivo* administration of locked-nucleic acid inhibitors of miR-22-3p (LNA-22) reduced lymphocyte accumulation in both the spleen and lymph nodes compared to LNA scramble (LNA-Scr) control-treated mice. Strikingly, LNA-22-3p treatments reduced kidney disease pathology and glomerular IgG deposition compared to LNA-Scr treatments in SLE mice. Moreover, miR-22-3p inhibition reduced the proportion of T effector memory IFN-γ producing CD4^+^ T cells, suggesting that miR-22-3p regulates Th1 T cell differentiation. We also found that miR-22 inhibition in mice reduced STAT1 phosphorylation in the kidney which was correlated with loss of IFN-γ production by splenic CD4^+^ T cells. In conclusion, our findings suggest that miR-22-3p is a critical regulator of SLE-associated CD4^+^ T cell immunity and kidney disease. These results provide therapeutic potential for limiting splenic Th1 signaling and preventing the progression of lupus nephritis.

**Key Findings:**

- Extracellular miR-22-3p levels are significantly increased in plasma from human SLE subjects.
- Inhibition of miR-22-3p *in vivo* significantly reduced lymphocyte accumulation in both the spleen and lymph nodes in a mouse model of SLE, thus reducing splenomegaly and lymphadenopathy.
- miR-22-3p inhibition significantly reduced IFN-γ expression and secretion from splenic T cell subsets.
- Inhibition of miR-22-3p *in vivo* resulted in decreased IgG deposition in the kidney, decreased STAT1 phosphorylation, and decreased kidney disease in a mouse model of SLE.

## Introduction

Systemic lupus erythematosus (SLE) is characterized by loss of tolerance to self, which leads to increased activation of lymphocytes and circulating autoantibodies[1]. The disease can affect nearly any organ, but nephritis is a major cause of morbidity and mortality in SLE patients[2]. The molecular mechanisms that link SLE to nephritis and other co-morbidities, e.g. cardiovascular disease (CVD), are not well-understood. As a result, therapeutic approaches are limited for SLE patients due to a paucity of druggable targets. Therefore, a greater understanding of immune dysregulation in SLE and lupus nephritis is needed for the development of effective treatments. Recently, RNA based therapies have emerged as a potential strategy to manipulate immune processes and treat inflammatory diseases[3–5]. One understudied potential regulatory mechanism in SLE and lupus nephritis is non-coding RNA-mediated gene regulation, particularly the biological relevance of extracellular microRNAs (miRNA).

miRNAs are post-transcriptional regulators of gene expression and have been demonstrated to regulate critical genes in lymphocyte phenotypes and SLE-associated pathways[6–8]. Recent studies have reported that altered miRNA activity contributes to T cell and B cell dysregulation in SLE. For example, miR-146a was found repress the pro-inflammatory AP-1 transcription factor complex, a key driver of interleukin-2 (IL-2) production and T cell activity[9], both of which are dysregulated in SLE. miR-24 was reported to directly target forkhead box P3 (FOXP3), a master transcription factor for regulatory T cells (Treg)[10]. Moreover, studies by Xiao *et al*. demonstrated that miRNAs in the miR-17-92 cluster promoted autoimmunity in mice through loss of activation-induced cell death in T cells[11]. A follow up study by Jiang *et al*. expanded on these findings and reported that the miR-17-92 cluster also supported IFN-γ production and antagonized Treg differentiation[12]. In addition to these miRNAs, many immune cell-associated miRNAs were also found to be altered in serum/plasma of SLE subjects and have been postulated as potential biomarkers for autoimmune disorders, including miR-223-3p, miR-92a-3p, and miR-20a-5p[13, 14]. These studies clearly demonstrate a role for miRNAs in SLE and lymphocyte biology; therefore, we aimed to extend the investigation of SLE miRNAs in the context of lupus nephritis which we posit will likely lead to a better understanding of the pathophysiology of SLE and co-morbidities.

Here, small RNA sequencing (sRNA-seq) was used to quantify miRNA changes in plasma from human SLE subjects, and miR-22-3p was identified as one of the most highly-abundant and differentially altered miRNAs in SLE plasma. Results from this study also demonstrate that inhibition of miR-22-3p using locked-nucleic acids (LNA-22) in a mouse model of SLE delayed the onset of disease activity and significantly decreased early anti-dsDNA antibody titer after 5 weeks compared to B6 control mice. Moreover LNA-22 treatments decreased activation and frequency of IFN-γ^+^ CD4^+^ T cells. Inhibition of miR-22-3p *in vivo* also reduced lupus nephritis and associated immune responses. Collectively, these data suggest that miR-22-3p likely contributes to T cell dysregulation in SLE and promotes lupus nephritis.

## Methods

### 1.1 Subjects

Clinical data and stored plasma from SLE (n=12) and healthy (n=12) subjects matched for age, race and sex were used for this study (Table S1). These subjects represent a subset of participants from two prior studies which were concurrently enrolled; study procedures were previously reported[15, 16]. Studies were conducted under approved Vanderbilt IRB protocols and written informed-consent was obtained from all subjects. Disease activity was measured in SLE subjects using the Systemic Lupus Erythematosus Disease Activity Index (SLEDAI), as modified for the SELENA trial[2]. Disease damage was assessed by the Systemic Lupus International Collaborating Clinics/American College of Rheumatology damage index (SLICC)[17].

### 1.2 Animals

B6.Cg-Foxp3^tm2Tch^/J mice (The Jackson Laboratory, strain #006772) were bred to B6.SLE1.2.3 mice originally obtained from Ward Wakeland (UTSW, Dallas, TX) and maintained in our colony (here after referred to as B6 and SLE, respectively). All mice were housed in a room with a 12h light:12h dark cycle with access to food and water *ad libitum*. Experiments involving mice were approved by the Vanderbilt Institutional Animal Care and Use Committee. SLE mice were retro-orbitally injected with 10 mg/kg of LNA-miR-22-3p (LNA-22) or scrambled control (LNA-Scr) (miRCURY LNA Inhibitor probe in-vivo, Qiagen) every 2 weeks beginning at 10-12 weeks of age, before the onset of disease (see Figure 2A). Animals were euthanized and analyzed 1 weeks after the final injection. Age-matched B6 mice functioned as untreated wild-type controls.

### 1.3 Lymphocyte Isolation

Splenic CD4^+^ T cells or CD19^+^ B cells were positively selected using immunomagnetic MicroBeads (Miltenyi Biotec) as per manufacturer’s instructions. Cells were washed and filtered through a LS magnetic column on a QuadroMACS separator (Miltenyi Biotec). Cells were then used in transcriptomic analysis. Alternatively, CD4^+^ T cells were stimulated with plate-bound α-CD3 and soluble α-CD28 antibodies (both at 2 μg/ml; Tonbo Biosciences) for 72 hours. Supernatants were collected and examined for cytokines by specific ELISA.

### 1.4 small RNA sequencing

To quantify miRNAs in human plasma, high-throughput small RNA sequencing (sRNA-seq) was performed[18, 19]. Briefly, total RNA was isolated from ethylenediaminetetraacetic acid (EDTA)-collected plasma using Total RNA Purification Kits (Norgen Biotek). Small RNA (cDNA) sequencing libraries were generated by TruSeq Small RNA Library Preparation Kits (Illumina). Libraries were size-selected using a Pippin Prep (Sage Science) and sequenced using the NextSeq500 platform (Illumina) at the Vanderbilt Technologies for Advanced Genomics (VANTAGE) DNA sequencing core facility. sRNA-seq data were analyzed using the TIGER (Tools for Integrative Genome Analysis of Extracellular sRNAs) pipeline[20]. miRNA aligned read counts were normalized to the total number of high-quality reads per sample and reported as Reads Per Million total reads (RPM). Differentially expression analysis for miRNAs was performed by DESeq2 with adjustment for batch effects[21]. Plasma miR-22-3p concentrations were validated in each sample by quantitative polymerase chain reaction (qPCR). For normalization of RNA extraction efficiency, plasma RNA samples were spiked with a standard of three exogenous single-stranded miRNA oligonucleotides (cel-miR-39, cel-miR-54, and cel-miR238; Qiagen) after the initial lysis step. A qScript microRNA cDNA synthesis kit (Quantabio) was used to prepare plasma cDNA and a miR-22-3p PCR assay (Quantabio) and PerfeCTa SYBR green supermix for iQ (Quantabio) were used for qPCR. Plasma miR-22-3p concentrations were derived from a standard dilution series of a known concentration of a DNA mimetic of the target miRNA sequence and normalized to the spike-in standards. To quantify miRNA expression in cells and tissues, total RNA was isolated from isolated splenic lymphocytes using Total RNA Purification kit (Norgen Biotek) and from tissues (e.g. kidney, spleen) using miRNeasy mini isolation kits (Qiagen).

### 1.5 Transcriptomics

To profile protein-coding gene expression, total RNA was isolated from splenic CD4^+^ T cells (isolated MACS) using the Total RNA Isolation kits (Norgen Biotek). Total RNA sequencing (RNA-seq) was performed with ribosomal RNA (rRNA)-depletion using the Ovation RNA-Seq Systems 1-16 for Model Organisms, mouse sequencing kit (NuGEN). Libraries were assessed for quality on the Bioanalyzer (Agilent) and quantified on the Qubit Fluorometer (Life Technologies) and then sequenced (paired-end 75 cycles) on the HiSeq3000 Illumina platform at the VANTAGE DNA sequencing core facility. Data analysis was performed using DEseq2For mRNA, reads were aligned to the GENCODE GRCm38.p5 genome using STAR v2.5.3a[22]. GENCODE vM16 gene annotations were provided to STAR to improve the accuracy of mapping. Quality control on raw reads was performed using FastQC (bioinformatics.babraham.ac.uk/projects/fastqc). FeatureCounts v1.15.2[23] was used to count the number of mapped reads to each gene. Significant differentially expressed protein coding genes with FDR-adjusted p-value < 0.05 and absolute fold change > 1.5 were detected by DESeq2 (v1.20)[21].

### 1.6 Flow cytometry

Splenocytes were incubated with Fc-Block (BD Biosciences) for 15 minutes at room temperature then stained for 30 minutes on ice with fluorophore conjugated antibodies diluted in FACS buffer containing HBSS, 1% BSA, 4.17 mM sodium bicarbonate, and 3.08 mM sodium azide. Antibodies are listed in Table S2. For intracellular cytokine staining, whole splenocytes were stimulated for five hours with PMA (20 ng/ml; Sigma) and ionomycin (1 μg/ml; Sigma) in the presence of GolgiStop (1.3 μg/ml; BD Biosciences) and stained using the FoxP3/Transcription Factor Staining Buffer Set (eBioscience). Briefly, samples were surface stained as described above, then fixed and permeabilized overnight at 4°C. The following day cells were stained for 30 minutes at room temperature with antibodies to intracellular antigens diluted in 1x permeabilization buffer. All samples were washed and resuspended in 2% PFA before acquisition on a seven-color MacsQuant flow cytometer (Miltenyi Biotec). Data were analyzed using FlowJo Single Cell Analysis software.

### 1.7 Kidney pathology scoring

Severity of glomerulonephritis was determined by an independent blinded pathologist at the Vanderbilt Translational Pathology Shared Resource core. A score of 0 indicated no pathology. A score of 1 represented a mild increase in basement membrane thickness and mild hypercellularity. A score of 2 signified moderate basement membrane and capillary loop thickening and moderate hypercellularity. A score of 3 indicated marked thickening of the basement membrane and capillary loops, marked hypercellularity, and necrosis. A score of 4 signified severe basement membrane thickening and capillary loop thickening, severe hypercellularity and inflammation, and obsolescence.

### 1.8 Antibody deposition in kidneys

Five-micron frozen kidney sections were incubated with goat-anti-Ig, anti-IgG_1_ and anti-IgG_2c_ diluted 1:50 in 5% normal goat serum for 1 hour at 37°C. Sections were washed and incubated with biotinylated rabbit anti-goat-Ig for 30 minutes (diluted 1:100) at 37°C. Sections were washed and incubated with either avidin-FITC or avidin-Texas Red and visualized by fluorescent microscopy.

### 1.9 ELISA

Anti-dsDNA IgG in mouse serum was measured as previously described[24]. 96-well Nunc MaxiSORP plates (Thermo Scientific) were pre-coated with methylated BSA (Sigma) diluted in 1x PBS (0.1 mg/ml) for 30 minutes at 37°C. Plates were then washed twice and coated with Calf Thymus dsDNA (Sigma) in 1x PBS (50 μg/ml) for 30 minutes at 37°C. Plates were washed twice and blocked overnight at 4°C in blocking buffer (3% BSA, 3mM EDTA, 0.1% gelatin in 1x PBS). After washing twice with 1x PBS, serum samples were applied diluted 1:1000 in serum diluent (2% BSA, 3mM EDTA, 0.05% Tween, and 0.1% gelatin in 1x PBS) and incubated for two hours at room temperature on an orbital shaker. Plates were washed twice with PBS-Tween followed by two washes with 1x PBS. IgG HRP (Promega) was applied to plates diluted 1:5000 in secondary diluent (1% BSA and 0.05% Tween in 1x PBS) and incubated overnight at 4°C. Plates were washed twice with PBS-Tween and twice with 1x PBS before being developed with OptEIA TMB Substrate (BD Biosciences). Reaction was quenched with 2N hydrochloric acid and plates were read at 450nm. IFN-γ ELISAs were performed according to manufacturer's instructions (BD Biosciences).

### 1.10 Western blotting

Whole kidney lysates were made by homogenizing tissue in RIPA buffer. Fifty μg of total protein was loaded on a 4-12% SDS-polyacrylamide gel under denaturing and reducing conditions. Proteins were transferred to nitrocellulose and subjected to Western blotting using rabbit anti-STAT1 antibodies (Cell Signaling) recognizing STAT1 (polyclonal) or pSTAT1^Tyr701^ (clone 58D6). Membranes were incubated with anti-rabbit IgG-IRDye 700nm (LiCor) and bands were visualized using the Odyssey system. Semi-quantitative analysis of Western Blots was conducted using the free software Image Studio Lite.

### 1.11 Statistics

Data are presented as mean± standard deviation or standard error of the mean (as noted in the text) for normally distributed data or median [interquartile range] for non-normally distributed data. Categorical data are presented as number (percentage). Significance was determined by Student’s *t*-test or Mann Whitney U non-parametric tests for continuous variables and Fisher exact test for categorical data. Two-sided *p*-values less than 0.05 were considered significant.

## 1 Results

### 2.1 miR-22-3p levels are increased in plasma from SLE subjects

To quantify changes in plasma miRNAs associated with SLE in human subjects, sRNA-seq was performed with RNA isolated from plasma collected from human SLE (n=12) subjects and age- and sex-matched healthy control (n=12) subjects. Remarkably, we found that plasma miRNAs (in total) were increased in plasma from SLE compared to control subjects, as 182 miRNAs were significantly increased and only 4 miRNAs were significantly decreased in SLE plasma compared to control plasma (Fig. 1A, Table 1). From the list of 182 significantly increased miRNAs, miR-22-3p was the most abundant, significantly differentially expressed miRNA in SLE plasma. To validate these sRNA-seq results, quantitative real-time PCR was used to quantify miR-22-3p concentration in plasma from SLE and control subjects, and we confirmed that miR-22-3p is significantly increased in SLE plasma (Fig. 1B). Previously, miR-22-3p was identified as one of three miRNAs that were found to be increased in circulating B cells isolated from SLE patients[25]. Therefore, extracellular miR-22-3p levels in circulation may represent increased miR-22-3p activity in cells and tissues.

**Figure 1.**
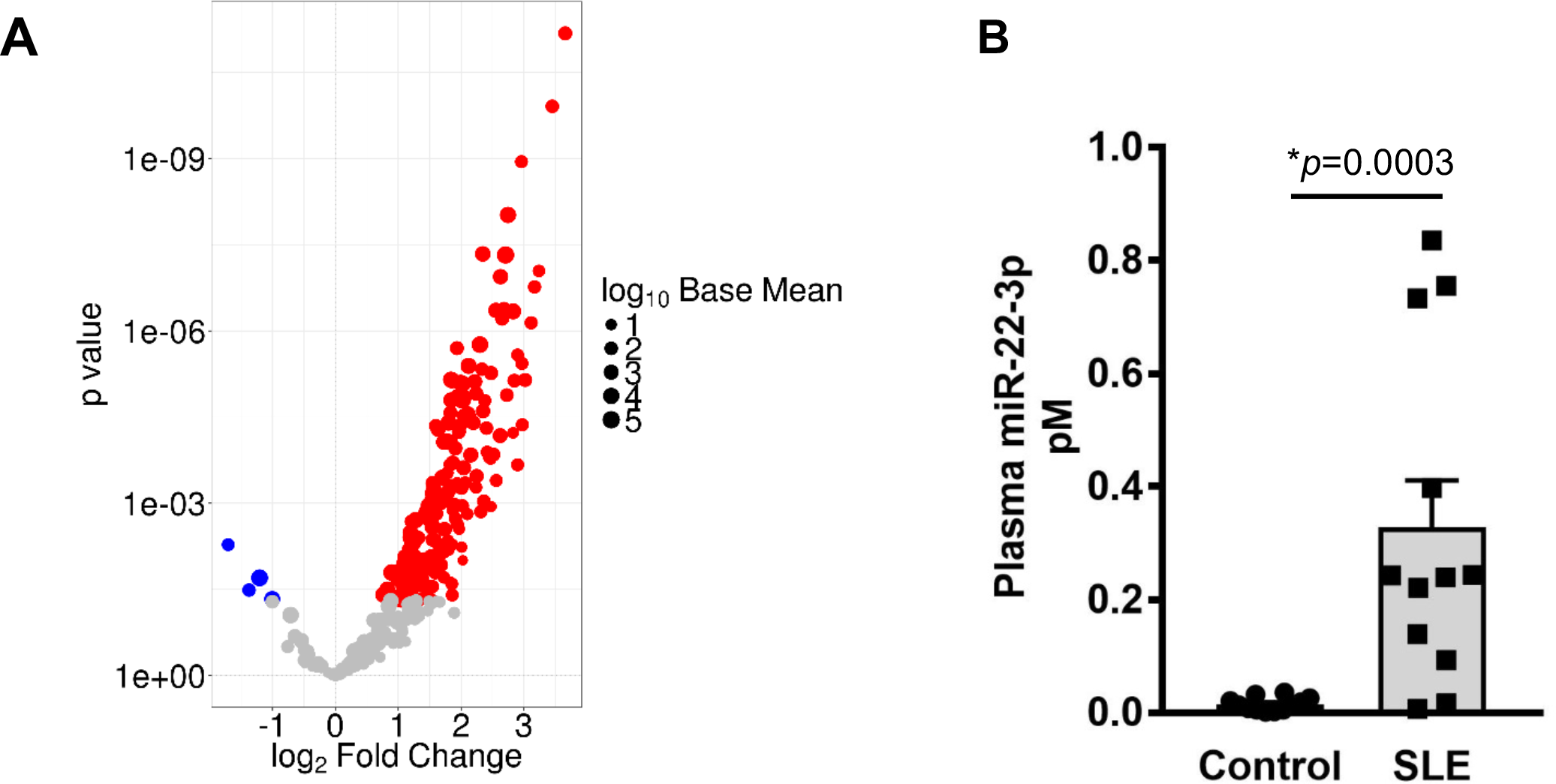
Circulating miR-22-3p levels are increased in plasma from human SLE subjects. **(A)** Significant differentially abundant miRNAs in plasma from SLE and healthy control subjects. Volcano plot. N=12. Red, significantly increased; blue, significantly decreased miRNAs. **(B)** qPCR of miR-22-3p levels in plasma from SLE and healthy control subjects. N=12, values are mean ± S.E.M. *p<0.05. Mann-Whitney non-parametric tests.

**Table 1.**
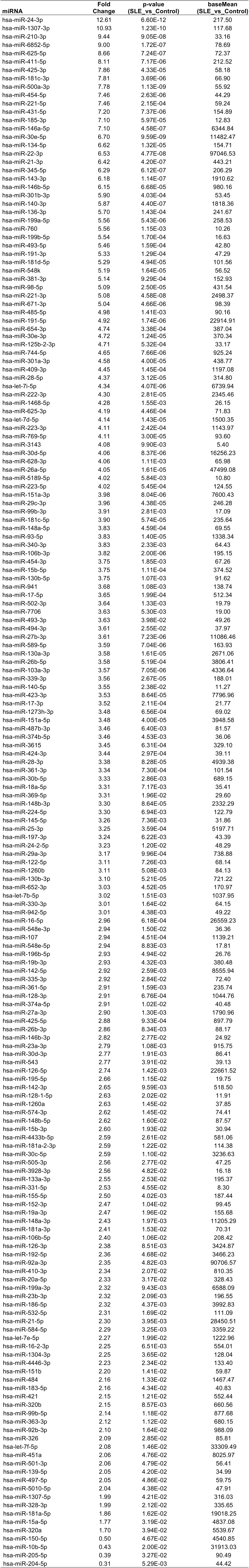
Significant differentially abundant plasma miRNAs in SLE.

### 2.2 miR-22-3p inhibition reduces splenomegaly and lymphadenopathy in a mouse model of SLE

Based on the high abundance of miR-22-3p in SLE patient plasma and patient B cells[25], we hypothesized miR-22-3p contributes to SLE pathogenesis and inhibition miR-22-3p may reduce autoimmunity and/or SLE-associated co-morbidities. To inhibit miR-22-3p *(in vivo)* in the setting of SLE, LNA inhibitors against miR-22-3p (LNA-22) or scrambled control (LNA-Scr) were injected intravenously (i.v.) into SLE mice. Briefly, prior to the onset of the disease, 10-12-week-old SLE mice were injected with 10 mg/kg of LNA-22 or 10 mg/kg LNA-Scr control (Fig. 2A). Mice were injected every 2 weeks for 10 weeks, and then sacrificed 1 week after the last injection. Cellular levels of miR-22-3p were significantly reduced in both splenic CD4^+^ T cells and CD19^+^ B cells in SLE mice treated with LNA-22 compared to LNA-Scr-treated, thus demonstrating that LNA-22 treatments suppressed cellular miR-22-3p levels in splenic lymphocytes (Fig. 2B and C). To demonstrate that LNA-22 treatments altered miR-22-3p activity in lymphocytes, real-time PCR was used to quantify the mRNA levels of an experimentally validated miR-22-3p target gene, phosphatase and tensin homolog (Pten)[26, 27]. In both splenic CD4^+^ T cells and CD19^+^ B cells, *Pten* mRNA levels were significantly increased in LNA-22-treated mice compared to LNA-Scr-treated SLE mice (Fig. 2D and E). To assess the systemic impact of miR-22-3p inhibition in the setting of SLE, relevant tissues were examined in SLE treated mice. Strikingly, LNA-22 treatments dramatically reduced the splenomegaly and lymphadenopathy normally associated with disease in SLE mice, as compared to LNA-Scr treatments (Fig. 2F). In response to miR-22-3p inhibition (LNA-22), spleen weights and cell numbers were also reduced compared to LNA-Scr treatments (Fig. 2G and H). There was not a significant difference in total body weight between LNA-22 and LNA-Scr-treated mice (data not shown). Collectively, these data suggest that miR-22-3p promotes splenomegaly and lymphadenopathy in SLE mice, and inhibition of miR-22-3p may be a viable strategy to limit the effects of SLE on the spleen and lymph nodes.

**Figure 2.**
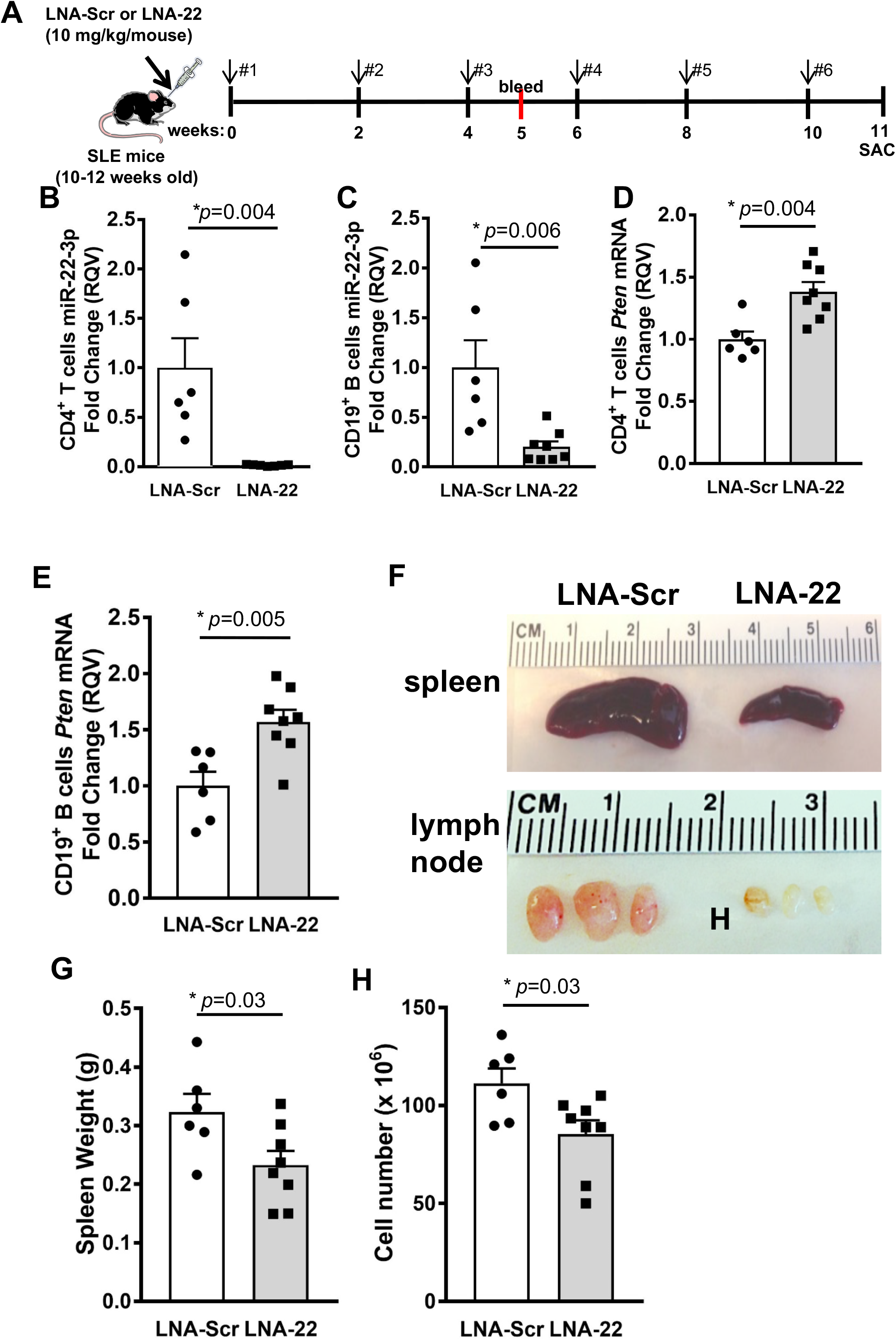
Inhibition of miR-22-3p reduces splenomegaly in SLE mice. **(A)** Schematic of experimental design for LNA injections. SLE mice were i.v. injected at the retro-orbital site with 10 mg/kg of either LNA-22 or LNA-Scr once every 2 weeks for 10 weeks, and mice were euthanized (SAC) 1 week following last injection. (B-E) Results from real-time PCR studies in splenic CD4^+^ T cells and CD19^+^ B cells. N=6-8. Unpaired Student's t-tests. **(B)** miR-22-3p levels in CD4^+^ T cells. **(C)** miR-22-3p levels in CD19^+^ B cells. **(D)** *Pten* mRNA levels in CD4^+^ T cells. **(E)** *Pten* mRNA levels in CD19^+^ B cells. **(F)** Representative spleens and lymph nodes from SLE mice treated with either LNA-22 or control LNA scrambled (LNA-Scr). **(G)** Spleen weights and total spleen cell numbers for SLE mice treated with LNA-22 or LNA-Scr. N=6-8. Values are mean ± S.E.M., *p<0.05. Unpaired Student's t-tests.

### 2.3 Inhibition of miR-22-3p reduces Th1 cells in SLE

Naïve T cells differentiate into specific T helper sub-sets, e.g. Th1, Th2, and Th17 cells and can later become T central (T_CEM_) or T effector memory (T_EM_) cells[28]. These different T cell phenotypes secrete characteristic cytokines and mediate distinct processes in immune responses to pathogens or host cells in the context of SLE[29]. In the spleen, LNA-22 treatments in SLE mice resulted in a significant increase in the percent naive CD4^+^ T cells (CD44^lo^CD62L^hi^) compared to LNA-Scr-treated mice (Fig. 3A and B). Moreover, we found a significant decrease in proportion of CD4^+^ T_EM_ cells (CD44^hi^CD62L^lo^) in LNA-22-treated compared to LNA-Scr treated SLE mice (Fig. 3A and C). This was accompanied by a significant increase in percentage of CD4^+^ T_CEM_ cells (CD44^hi^CD62L^hi^) following LNA-22 treatments (Fig. 3A and D). To determine the impact of miR-22-3p inhibition on cytokine expression, intracellular flow cytometry was conducted, and we found a significant reduction in the percentage of IFN-γ^+^ CD4^+^ T cells in LNA-22-treated compared to LNA-Scr-treated mice (Fig. 3E and F). Consistently, supernatants from activated splenic CD4^+^ T cells (anti-CD3/CD28 stimulation for 72h) from LNA-22-treated mice contained significantly less IFN-γ compared to CD4^+^ T cells from LNA-Scr-treated B6.SLE mice (Fig. 3G). Of note, we failed to find differences in IL-17 and IL-4 production by CD4^+^ T cells (data not shown). These data suggest that treatment of SLE mice with LNA-22 decreases the activation and differentiation of IFN-γ^+^ CD4^+^ T cells. Overall, these results suggest that miR-22-3p promotes Th1 T cell responses in SLE which can be readily targeted *in vivo* using LNA-22.

**Figure 3.**
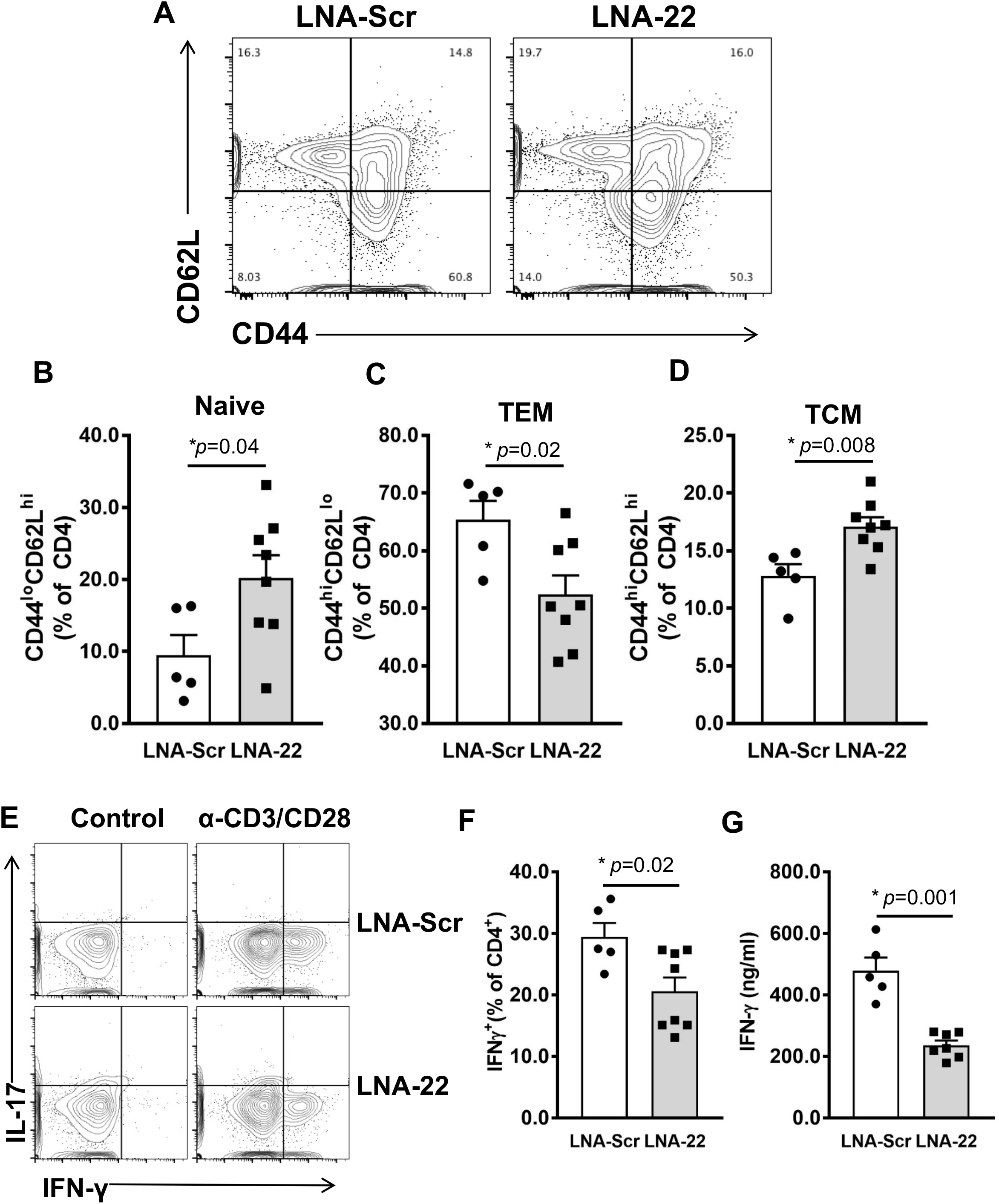
Inhibition of miR-22-3p reduces T cell activation and Th1 responses in SLE mice. Splenic CD4^+^ T cells from SLE mice treated with LNA-22 or LNA-Scr as described in Fig 1 were analyzed for activation phenotypes via expression of CD44 and CD62L by flow cytometry. **(A)** Representative flow plots of splenic CD4^+^ T cells from SLE mice treated with LNA-22 or LNA-Scr. (B-C) Quantification of percent CD4^+^ T cells expressing CD44 and CD62L. N=5-8. **(B)** Naïve T cells. **(C)** T_EM_. **(D)** T_CM_. **(E)** Intracellular flow cytometry analysis of interleukin 17 (IL-17) and interferon gamma (IFN-γ) in splenic CD4^+^ T cells isolated from SLE mice treated with either LNA-Scr (top panels) or LNA-22 (bottom panels). **(F)** Percentage of CD4^+^ T cells producing IFN-γ as determined by intracellular flow cytometry. N=5-8. **(G)** Secreted IFN-γ in media of splenic CD4^+^ T cells following stimulation with anti-CD3 and anti-CD28. N=5-7. Values are mean ± SEM, *p<0.05. Unpaired Student’s t-tests.

### 2.4 Inhibition of miR-22-3p significantly alters T cell gene expression in SLE mice

To determine the impact of miR-22-3p inhibition on gene expression in SLE mice, CD4^+^ T cells were isolated by magnetic-activated cell sorting (Miltenyi Biotec) from spleens of mice treated with LNA-22 or LNA-Scr control. Protein coding gene (mRNA) expression was quantified in sorted CD4^+^ T cells by total RNA sequencing (RNA-seq). Differential expression analyses were completed using DEseq2 with a Benjamini-Hochberg FDR-corrected (adjusted) p<0.05 and absolute fold change >1.5 considered a significant change[21]. Strikingly, LNA-22 treatments in mice resulted in a significant change to 115 genes (mRNA) in splenic T cells, as compared to LNA-Scr controls (Fig. 4A, Table S3). From this list of 115 altered genes, 69 genes were found to be significantly increased and 46 genes to be significantly decreased in CD4^+^ T cells (Fig. 4A, Table S3). Furthermore, we identified 10 putative target genes for miR-22-3p within the list of the 69 significantly increased genes (mRNAs), suggesting that LNA-22 inhibition of miR-22-3p likely reduced miR-22-3p activity in mouse splenic T cells (Table S3). Though most gene (mRNA) expression changes associated with LNA-22 treatments were not predicted to be the result of direct miR-22-3p regulation, we do not believe this diminishes the value of the data as most miRNA-regulated genes are likely affected through indirect mechanisms. Therefore, to assess the potential impact of these gene expression changes, pathway analyses were performed using MetaCore software (Agilent) which revealed a significant enrichment of altered genes associated with T cell regulatory pathways (Table S4), e.g. immune response T cell subsets for cell surface markers (Fig. S2) and secreted signals (Fig. S3). Nevertheless, the numbers of genes (objects in pathways) that were significantly altered in these enriched pathways were low despite the statistical significance of the pathway overall. Based on these pathways and prior associations with T cells, autoimmunity, or general immune response, candidate genes were selected for down-stream validation by real-time PCR, including C-X-C chemokine receptor type 3 (*Cxcr3*) and interleukin 21 (*Il21*) which were found to be significantly decreased by RNA-seq analysis (Table S3). To validate these changes, real-time PCR was used to quantify mRNA levels in splenic CD4^+^ T cells isolated from LNA-22 and LNA-Scr-treated mice, and we confirmed that of *Cxcr3* and *Il21* mRNA levels were significantly decreased in T cells *in vivo* with LNA-22 treatments (Fig. 4B and C). These data suggest that inhibition of miR-22-3p *in vivo* suppressed pro-inflammatory gene expression in CD4^+^ T cells in a mouse model of SLE. Moreover, these results support that miR-22-3p likely contributes to the pathophysiology of SLE.

**Figure 4.**
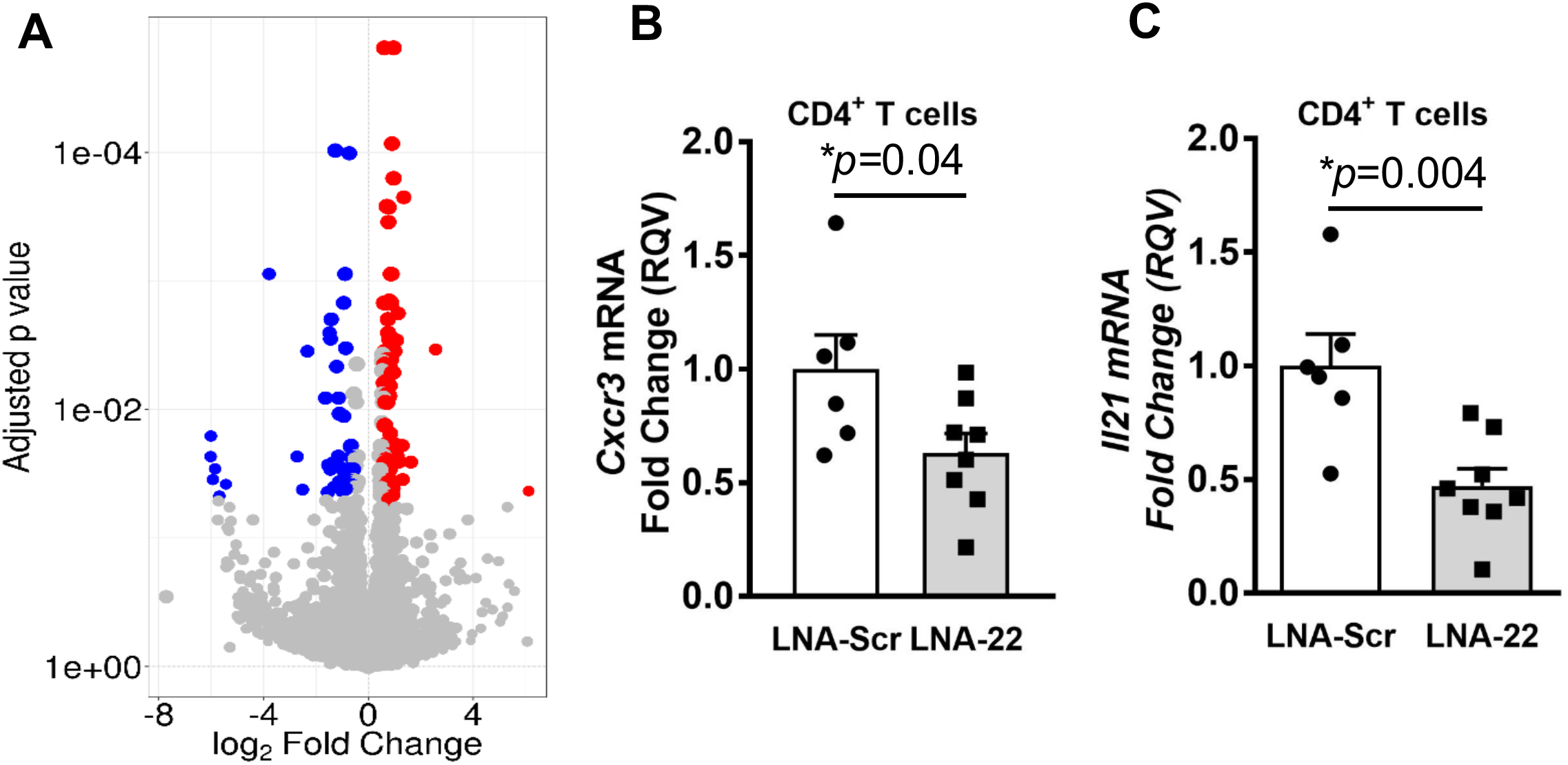
Inhibition of miR-22-3p results in significantly altered gene (mRNA) expression in splenic T cells isolated from SLE mice. **(A)** Significantly altered protein coding gene (mRNA) expression in CD4^+^ T cells isolated from SLE mice treated with LNA-22 or LNA-Scr. Volcano plot. Red, significantly increased; blue, significantly decreased genes (mRNA). Benjamini-Hochberg adjusted FDR p-values vs. log2 fold changes. (B-C) Real-time PCR of candidate genes in splenic CD4^+^ T cells. Relative quantitative values (RQV) reported as fold changes. N=5-8, values are mean ± S.E.M., *p<0.05. Unpaired Student’s t-tests. **(B)** *Cxcr3* mRNA levels. **(C)** *ll21* mRNA levels.

### 2.5 Inhibition of miR-22-3p reduces dsDNA titers

Production of autoantibodies such as anti-dsDNA titers are a principle biomarker for lupus activity[30]. Therefore, serum antibodies against dsDNA were measured in SLE mice treated with LNA-22 or LNA-Scr controls at baseline (10-12 weeks old), in the middle of the study (5 weeks treatment, mice 15-17 weeks old), and at the end of the study (11 weeks treatment, mice 21-23 weeks old). Interestingly, after 5 weeks of treatment, the LNA-22-treated mice were found to have significantly lower anti-dsDNA antibody titers compared to mice treated with LNA-Scr control (Fig. 5A). This difference was not observed, however, at the end of the study when mice were 21-23 weeks of age, as LNA-22-treated mice had reduced but not statistically significant differences in anti-dsDNA antibody titers. This observation was reflected in both anti-dsDNA IgG_1_ and IgG_2c_ levels (Fig. S4A and B). Moreover, we failed to find a significant difference in plasma cell proportions at the end of the study (Fig. S1). Therefore, we conclude that although LNA-22 treatments can delay the onset of SLE as defined by anti-dsDNA antibody production, it does not completely prevent the break in B cell tolerance in our mouse model of SLE.

**Figure 5.**
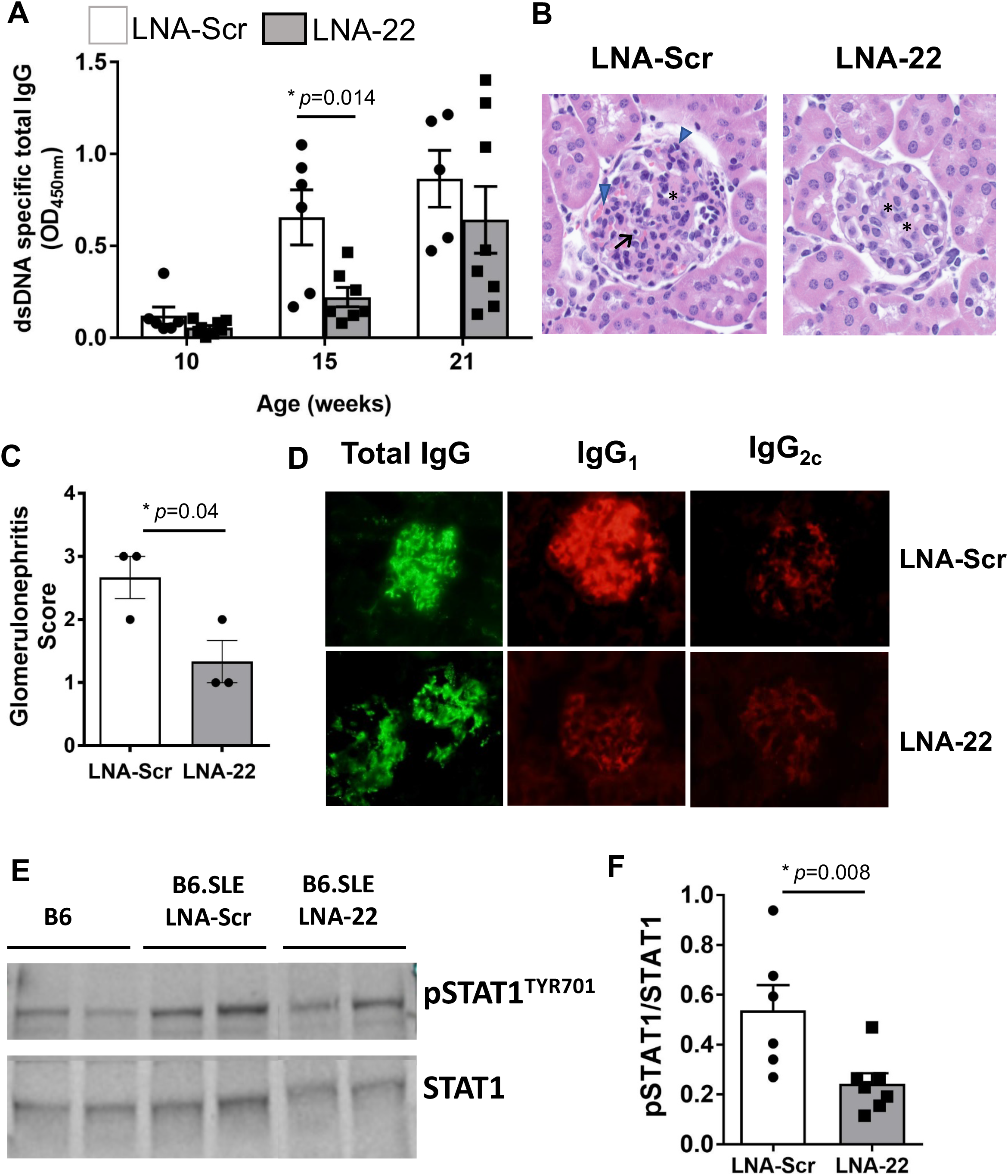
Inhibition of miR-22-3p reduces lupus nephritis in mice. **(A)** Serum levels of anti-dsDNA auto-antibodies measured by ELISA at baseline (10-12 weeks of age), post-5 weeks of treatments (15 weeks of age), and at the end of the study (21 weeks of age). N=5-8. **(B)** H&E staining of kidney paraffin sections from SLE mice treated with LNA-22 or LNA-Scr. H&E staining. **(C)** Glomerulonephritis scores. N=3. **(D)** Total IgG, IgG_1_, and IgG_2c_ deposition in kidney glomeruli of SLE mice treated with LNA-22 or LNA-Scr. **(E)** Western blotting of STAT1 phosphorylation (pSTAT1^TYR701^) in whole kidney lysates of treated SLE mice and B6 control mice. **(F)** Ratio of pSTAT1^TYR701^/STAT. N=6-7. Values are mean ± S.E.M., *p<0.05. Unpaired Student’s *t*-test.

### 2.6 miR-22-3p inhibition reduces SLE-associated kidney disease

Renal damage associated with lupus nephritis is one of the most detrimental co-morbidities of SLE[31]. Lupus nephritis is characterized by an influx of immune cells to the kidney, specifically, Th1 cell infiltration of the glomeruli[32]. To determine if miR-22-3p contributes to SLE-associated kidney disease and if inhibition of miR-22-3p improves renal pathology in SLE, kidneys were collected from SLE mice treated with LNA-22 or LNA-Scr controls. Based on H&E staining of kidney sections, LNA-22 treatments dramatically reduced glomerulonephritis severity in SLE mice, as compared to LNA-Scr treatments (Fig. 5B and C). Specifically, almost all glomeruli were diffusely affected, showing increased cellularity, increased eosinophilic hyaline material expanding the glomerular tuft, thickening of capillary loops, and occasional presence of single cell death. Kidneys from LNA-22-treated SLE mice showed significantly decreased disease severity (mean score of 1.5=mild glomerulonephritis) with the few glomeruli that were affected having a mild segmental increase in eosinophilic hyaline material in the glomerular tuft (Fig. 5B and C). To determine if the observed reduction in glomerulonephritis severity was linked to reduced immunoglobin (IgG) deposition in the kidney, fluorescence-immunohistochemistry was used to assess IgG in the glomeruli. Strikingly, we found that total IgG, as well as IgG_1_ and IgG_2c_ isotypes, were decreased in glomeruli of LNA-22-treated SLE mice compared to age-matched LNA-Scr-treated mice (Fig. 5D). Collectively, these data demonstrate that inhibition of miR-22-3p *in vivo* significantly decreased kidney pathology, specifically glomerulonephritis, in a mouse model of SLE.

To identify a potential mechanism that may underlie the molecular link between miR-22-3p and lupus nephritis, IFN-γ signaling was assessed in kidneys since LNA-22 treatments reduced IFN-γ expression in T cells and IFN-γ signaling is associated with lupus-associated kidney pathology[33]. To determine whether LNA-22 treatment resulted in reduced IFN-γ signaling, STAT1 activation was assessed by immuno-blotting phosphorylated STAT1 (pSTAT1) in whole kidney lysates. Interestingly, pSTAT1 levels were found to be significantly decreased in kidneys of LNA-22 treated mice compared to LNA-Scr control-treated SLE mice, when normalized to total STAT1 protein levels (Fig. 5E and F). Furthermore, the levels of pSTAT1 positively correlated to IFN-γ secretion in splenic T cells (Fig. S5). These data suggest that miR-22-3p regulation of IFN-γ production by CD4^+^ T cells may contribute to SLE and associated comorbidities, e.g. kidney disease.

## 3 Discussion

Extracellular miRNAs are a new class of biomarkers for autoimmunity, however, they may also contribute to SLE disease pathogenesis and lupus nephritis. Overall, results support that miR-22-3p contributes to SLE-associated nephritis, as inhibition of miR-22-3p *in vivo*, delayed or significantly reduced many immunologic features of SLE and lupus nephritis. Here, we report that plasma levels of miR-22-3p levels are significantly increased in human subjects with SLE. Although a link between miR-22-3p and lupus nephritis has not been previously reported, miR-22-3p has been linked to SLE and other chronic inflammatory diseases. For example, miR-22-3p has been reported to be elevated in B cells from SLE subjects compared to healthy human subjects[25]. Additionally, Pei *et al*. recently reported that miR-22-3p levels are elevated in peripheral CD4^+^ T cells in inflammatory bowel disease [34].

To determine the functional relevance of miR-22-3p in SLE, LNA inhibitors against miR-22-3p were i.v. injected into a mouse model of SLE every 2 weeks over 11 weeks. Moreover, LNA-22-treated SLE mice also showed decreased splenomegaly and lymphadenopathy compared to SLE mice treated with LNA-Scr controls. Although the impact of miR-22-3p inhibition on spleen size has not been previously reported, LNA inhibition of another miRNA (miR-21-5p) was also previously found to reduce splenomegaly in lupus[35]. Of note, we found that miR-21-5p levels, like miR-22-3p, were significantly increased in plasma from human SLE subjects; however, miR-21-5p levels were not as abundant as miR-22-3p. In addition to anatomical changes and cell counts, miR-22-3p inhibition *in vivo* was found to cause a significant change to splenic T cell gene expression, as we identified 115 genes (mRNA) that were significantly altered >1.5-fold after FDR (p-value) correction. Many of these gene changes are likely the result of indirect gene regulatory modules for miR-22-3p, as only a few putative miR-22-3p targets genes were found to be significantly increased with LNA-22 treatments in SLE mice. This observation is not uncommon and a majority of miRNA regulatory networks in all cells are likely through indirect mechanisms. These findings strongly support that miR-22-3p contributes to T cell gene regulation in SLE which can be targeted through LNA-based inhibition of miR-22-3p *in vivo*.

Anti-dsDNA antibody titers are a hallmark for SLE disease activity, and thus, we quantified the levels of anti-dsDNA titers in serum from LNA-22-treated mice at baseline, in the middle of the study (5 weeks), and at the conclusion (11 weeks). Interestingly, miR-22-3p inhibition significantly decreased anti-dsDNA antibody titers at the early (5 week) time-point compared to the LNA-Scr treatments; however, at the terminal time-point, no significant differences in anti-dsDNA titers were found between the treatments. These results suggest that miR-22-3p inhibition likely delays the onset of SLE through changes to early autoreactive B cells but does not fully inhibit the break in B cell tolerance. This observed response to miR-22-3p inhibition is consistent with the targeting of other miRNAs. For example, inhibition of miR-21 also dramatically reduced splenomegaly and lymphocyte proliferation but failed to affect anti-dsDNA titers at 12 weeks of treatments[35]. Moreover, over-expression of miR-326 was found to increase B cell auto-antibody production within 1 week[36]. Therefore, results from our studies and others suggest that anti-dsDNA auto-antibody production is phasic and miR-22-3p, along with other miRNAs, contribute to early titer production.

One of the most important observations from this study was that inhibition of miR-22-3p *in vivo* significantly reduced glomerulonephritis in our mouse model of SLE. Lupus nephritis is one of the most common and severe complications in SLE and a critically important predictor of morbidity and mortality in SLE. Lupus nephritis is characterized by an increase in Th1 signaling and deposition of immune complexes[37]. In our study, histology was used to assess renal pathology, and LNA-22 treatments were found to significantly reduce glomerular capsule size. Blinded pathological scoring confirmed these observations and results support the model in which miR-22-3p inhibition reduces nephritis in SLE mice. These findings were also accompanied by a reduction in immune IgG complex deposition in the kidneys, a hallmark of glomerular inflammation and lupus nephritis[37]. Reducing immune complex deposition is vital to preventing and treating lupus nephritis, as localization of immune complexes within the glomeruli can lead to complement activation and complement-mediated damage[38]. Moreover, immune complex deposition in the kidney promotes recruitment and activation of neutrophils and myeloid cells via their Fc-receptors [39, 40]. Activated immune cells secrete chemokines and cytokines which recruit additional immune cells that further exacerbate glomerular damage and lead to a pro-inflammatory environment in the kidney.

Due to the observed reduction in markers of SLE by LNA-22 treatments, we sought to determine if this was due to a reduction in immune cell activation. During lupus, T cell abnormalities include T cell hyperactivation, accumulation of effector memory T cells (CD44^hi^CD62L^lo^), as well as increased IFN-γ production[41]. T_EM_ cells, which can stimulate cytokine production rapidly following migration to inflamed tissues in SLE[42], were significantly reduced following LNA-22 treatment in SLE mice. This is in contrast to TEM cells (CD44^hi^CD62L^hi^) that have minimal cytokine-secreting capacity and were found elevated following LNA-22 treatments. Inhibition of miR-22-3p also reduced the proportion of IFN-γ positive CD4^+^ T cells as well as the secretion of IFN-γ following stimulation *ex vivo*. Classically, Th1 producing IFN-γ cells were postulated to promote cell-mediated immune diseases while their counterpart Th2 cells, which produce IL-4, drive autoantibody-mediated autoimmune disease. Th1 IFN-γ producing cells are common in autoimmune lupus disease[43]. Indeed, increased IFN-γ is observed in several lupus mouse models, particularly in the MRL-*Fas^lpr^* strain, and is associated with immune complex deposition in kidneys[44]. Several other miRNAs have previously been shown to regulate Th1 and IFN-γ signaling, including the miR-17-92 cluster[12], miR-125b[45], and miR-29[46]; however, in this study, we present a novel role for miR-22-3p in the Th1 population during SLE and lupus nephritis.

IFN-γ is a critical effector molecule in the progression of SLE and associated co-morbidities[43]. We found that the link between miR-22-3p and renal disease in lupus may be through IFN-γ mediated pSTAT1 activation. In our study, we found that STAT1 signaling, as reported by pSTAT1^Tyr721^ phosphorylation levels normalized to total STAT1 protein levels, was reduced in kidneys in SLE mice treated with LNA-22 compared to LNA-Scr. Moreover, pSTAT1 levels positively correlated with splenic CD4^+^ T cell IFN-γ secretion. Reduction of pSTAT1 decreases STAT1 transcriptional activity and leads to the down-regulation of IFN-inducible genes[47]. For instance, pSTAT1 has been shown to transactivate T-box transcription factor 21 (*Tbx21*), and Tbx21 drives IFN-γ production[48, 49]. Therefore, miR-22-3p may induce loss of pSTAT1 signaling thus contributing to reduced IFN-γ expression in the kidney through this novel gene regulatory module and feedback mechanism.

Collectively, the data suggest that miR-22-3p has great potential as a novel drug target to reduce glomerulonephritis and SLE pathophysiology. Although the full potential and viability of miR-22-3p as a drug target in SLE remains to be seen, particularly with regard to diseases like cancer, anti-miR-22-3p strategies were found in a separate study to suppress Th17 activity and inflammation in mice[50]. Moreover, in emphysema, inhibition of miR-22-3p via intranasal administration of LNA reduced inflammation and suppressed Th17 activity in mice, and *Mir22^-/-^* mice were found to have fewer Th17 cells in the lungs after cigarette smoke exposure and less Th17-mediated inflammation[50]. These previous reports support that low miR-22-3p expression in lymphocytes is critical to maintaining normal immune homeostasis

## Conclusions

Results from this study support that miR-22-3p is increased in plasma of SLE patients compared to matched healthy controls. This is supported by *in vivo* data demonstrating that inhibition of miR-22-3p in SLE mice reduces splenomegaly and T cell infiltration in tissues, alters T cell gene expression, delays the onset of lupus and anti-dsDNA titers, and decreases renal disease. Therefore, miR-22-3p holds great potential as a viable candidate for further investigation as biomarker or a drug target to prevent and treat SLE and lupus nephritis.

## Acknowledgements

We would like to acknowledge Kelli Boyd, DVM, PhD, DACVP for scoring kidney sections as well as the Translational Pathology Shared Resource supported by NCI/NIH Cancer Center Support Grant 2P30 CA068485-14 and the Vanderbilt Mouse Metabolic Phenotyping Center Grant 5U24DK059637-13.

## Supplementary data

Please find supplementary materials online.

