## Supplemental materials for "Inhibition of miR-22-3p reduces kidney disease associated with systemic lupus erythematosus"

**Supplementary Materials**

**Supplementary Figures.**

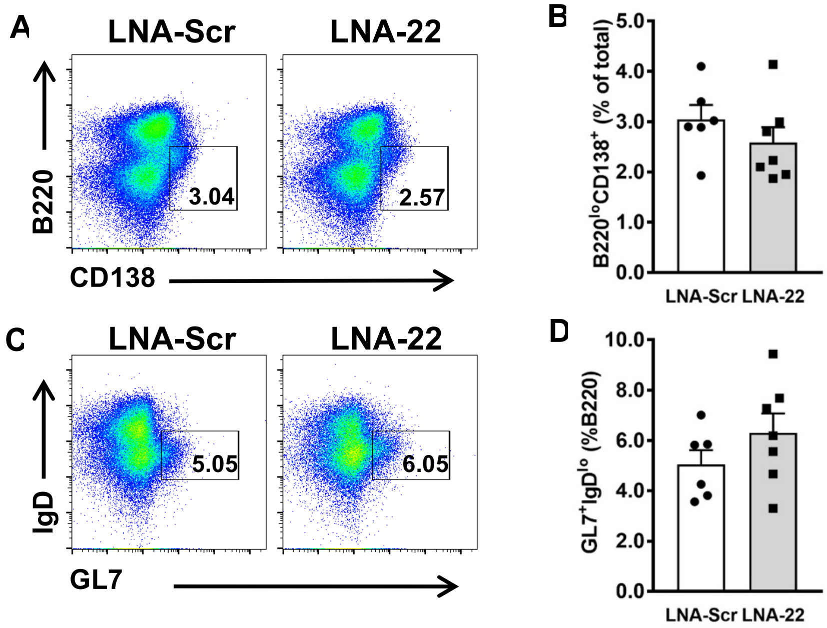


**Figure S1. Inhibition of miR-22-3p did not alter B cell subsets. (A)** Representative flow plots of splenic plasma B cells from SLE mice treated with LNA-22 or LNA-Scr. **(B)** Quantification of % total splenocytes expressing B220$^{lo}$ and CD138. N=6-7. **(C)** Representative flow plots of splenic germinal center B cells from SLE mice treated with LNA-22 or LNA-Scr. **(B)** Quantification of % total splenocytes expressing GL7 and IgD. N=6-7. Values are mean ± S.E.M.

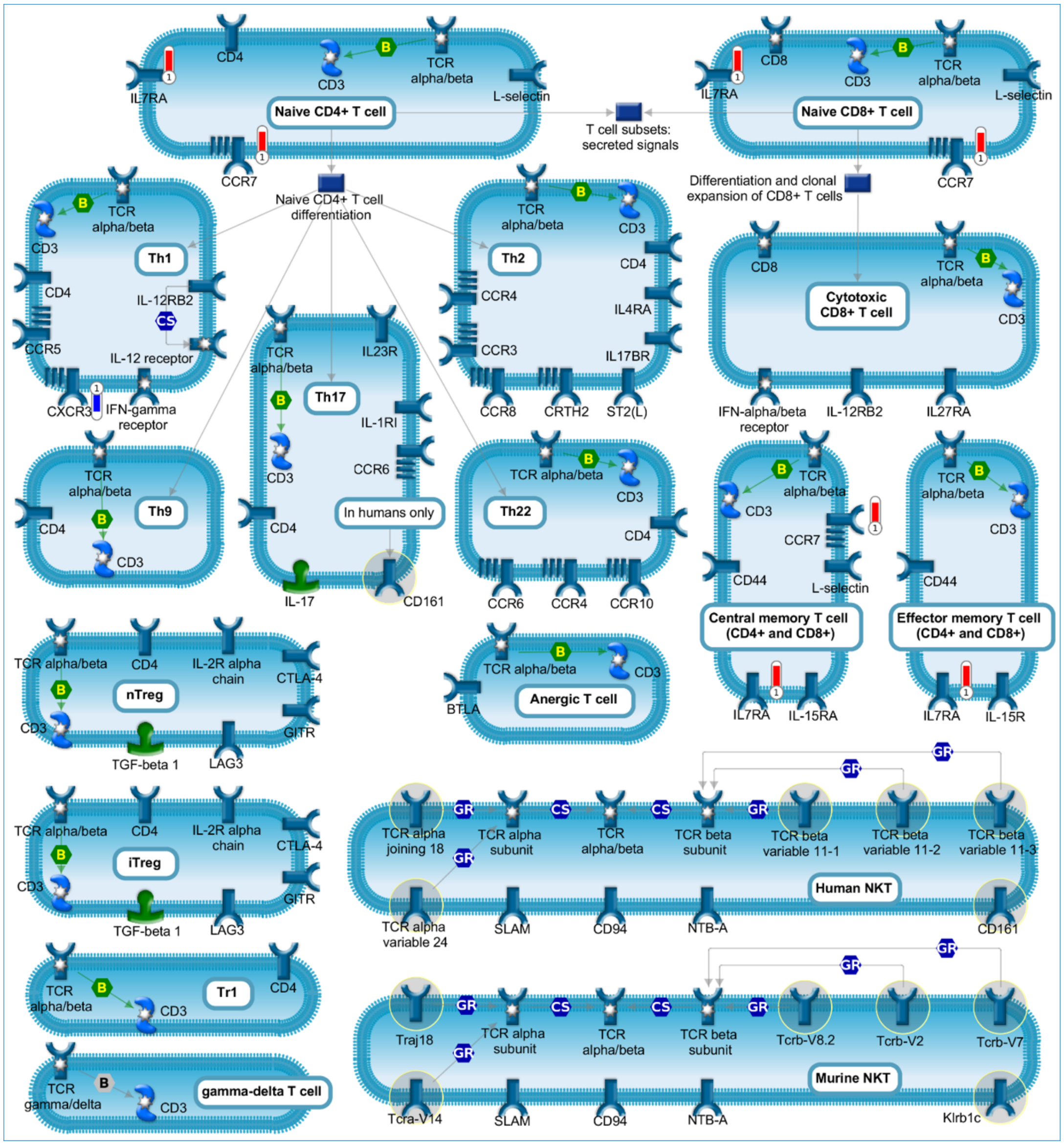


**Figure S2. Pathway analyses of T cell regulatory pathways.** RNA-seq pathway analyses performed on splenic $CD4^+$ T cells isolated from SLE mice treated with LNA-22 or LNA-Scr using MetaCore software (Agilent).

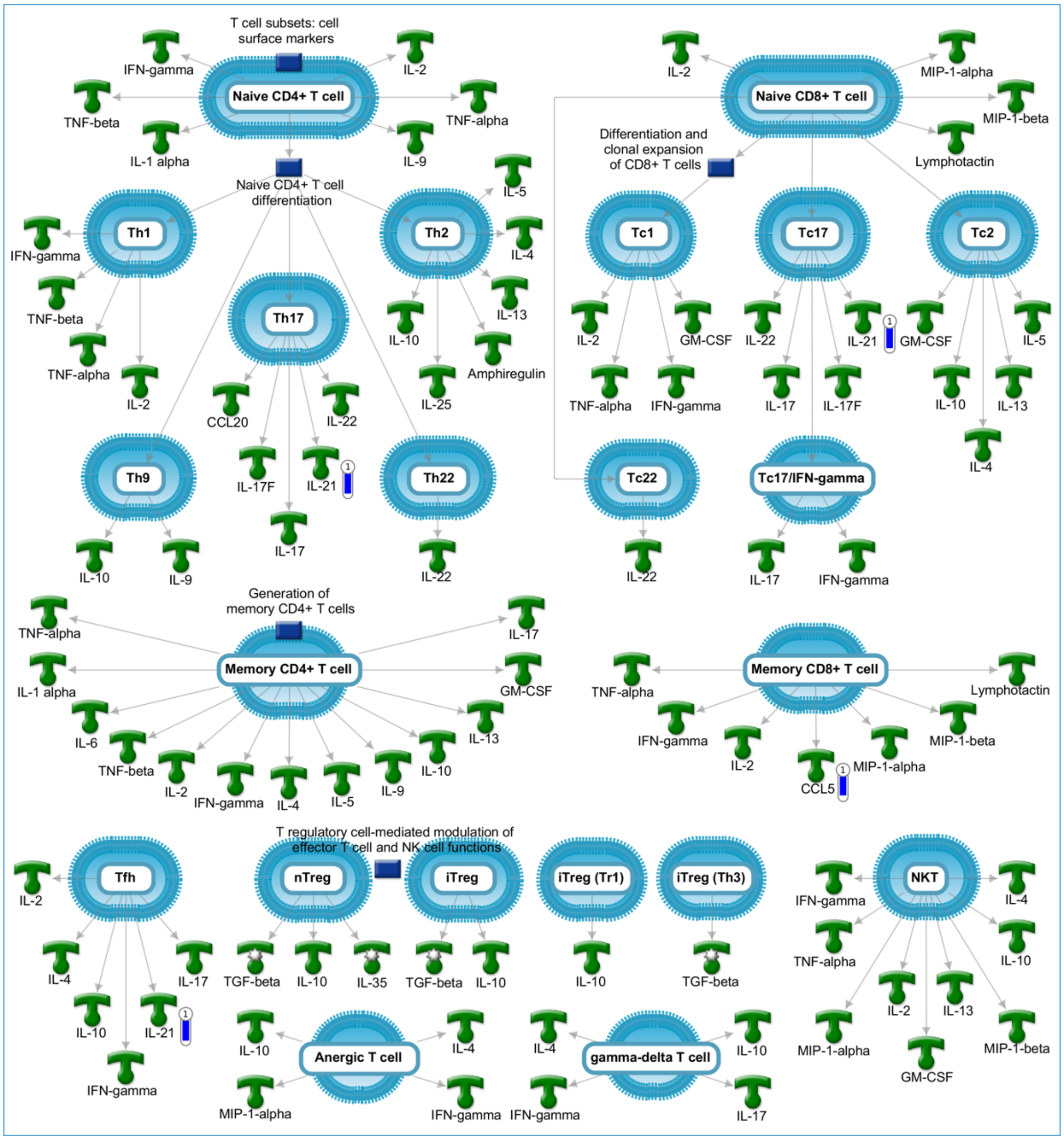


**Figure S3. Pathway analyses of T cell surface markers.** RNA-seq pathway analyses performed on splenic CD4$^+$ T cells isolated from SLE mice treated with LNA-22 or LNA-Scr using MetaCore software (Agilent).

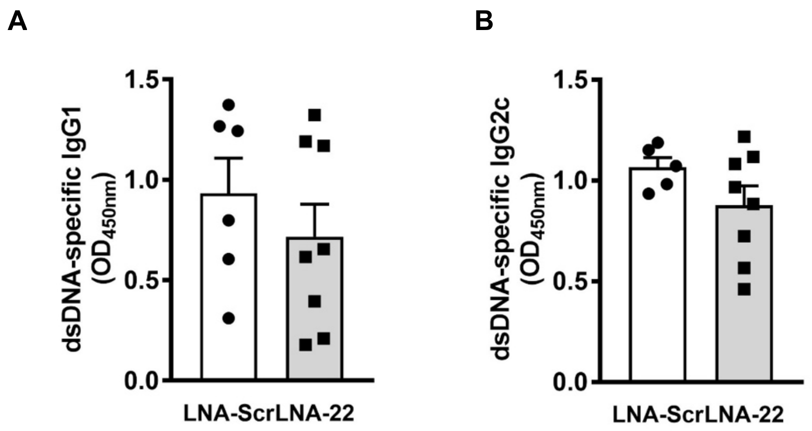


**Figure S4. Anti-dsDNA autoantibody titers of $IgG_1$ and $IgG_{2c}$.** Serum levels of anti-dsDNA autoantibodies $IgG_1$ **(A)** and $IgG_{2C}$ **(B)** measured by ELISA at the end of the study (21 weeks of age). N=5-8. Values are mean are mean ± S.E.M.

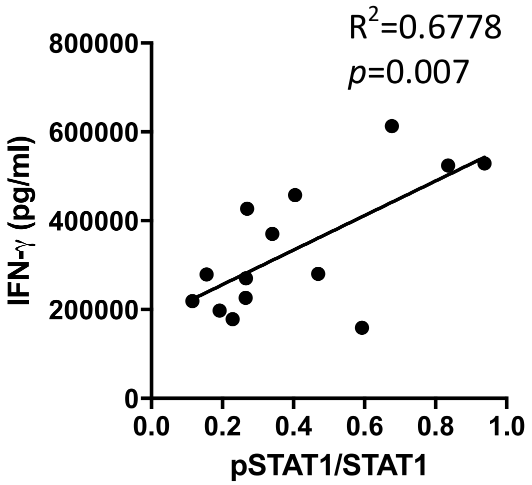


**Figure S5. Correlation CD4$^+$ IFN-γ production and kidney pSTAT1.** Correlation between IFN-γ production of splenic CD4$^+$ T cells and pSTAT1 levels in the kidney.

**Supplementary Tables.**

**Table S1. Subject characteristics**

| | SLE (N=12) | Control (N=12) | P |
|---|---|---|---|
| Age, years | 48 ± 11 | 49 ±11 | 0.77 |
| Race, #Caucasian | 12 (100) | 12 (100) | 0.99 |
| Sex, #female | 7 (58) | 7 (58) | 0.99 |
| Creatinine, mg/dl | 1.0 ±0.33 | 0.9 ±0.3 | 0.46 |
| SLEDAI score, units | 4 [0, 6] | - | - |
| SLICC score, units | 1 [0, 3] | - | - |
| Lupus nephritis (ever), # | 1 (8) | - | - |
| Hydroxychloroquine, # | 7 (58) | - | - |
| Methotrexate, # | 1 (8) | - | - |
| Mycophenolate mofetil, # | 1 (8) | - | - |
| Azathioprine, # | 2 (17) | - | - |
| Prednisone, # | 6 (50) | - | - |
| miR-22-3p, Ct | 27.0 ± 2.0 | 30.7 ± 1.4 | 0.0003 |
| miR-22-3p, pM (3- spike) | 0.33 ± 0.29 | 0.015±0.012 | 0.0003 |

Continuous data are presented as mean ± standard deviation. Categorical data are presented as number (percentage). Difference was determined by Mann Whitney U for continuous and Chi square for categorical data. Medications are listed as current use.

**Table S2. Antibodies for flow cytometry**

| Antigen | Clone | Manufacturer |
|---|---|---|
| CD69 | H1.2F3 | BD Biosciences |
| CD19 | 1D3 | Tonbo Biosciences |
| CD11b | M1/70 | Tonbo Biosciences |
| TCR-β | H57-597 | Tonbo Biosciences |
| CD8α | 53-6.7 | BD Biosciences |
| CD62L | MEL-14 | BD Biosciences |
| CD1d | 1B1 | BD Biosciences |
| CD40 | 1C10 | eBioscience |
| PD-1 | J43.1 | Tonbo Biosciences |
| MHC-II | AF6-120.1 | eBioscience |
| CD4 | GK1.5 | Tonbo Biosciences |
| CD23 | B3B4 | BD Biosciences |
| CD11c | N418 | eBioscience |
| CXCR5 | SPRCL5 | eBioscience |
| CD25 | PC61.5 | Tonbo Biosciences |
| CD5 | 53-7.3 | BD Biosciences |
| CD86 | GL1 | BD Biosciences |
| CD44 | IM7 | Tonbo Biosciences |
| B220 | RA3-6B2 | Tonbo Biosciences |
| F4/80 | BM8 | Tonbo Biosciences |
| IL-17a | TC11-18H10 | BD Biosciences |
| IFN-γ | XMG1.2 | Tonbo Biosciences |
| IL-10 | JES5-16E3 | BD Biosciences |

**Table S3. Significant differentially expressed genes (mRNA) in splenic CD4$^{+}$ T cells from mice treated with LNA-22.**

| GeneSymbol | Fold Change | padj | miR-22-3p target |
|---|---|---|---|
| Slitrk2 | 69.56110286 | 0.04315034 | |
| Cd36 | 5.937081808 | 0.003417657 | Yes |
| Rpl10-ps3 | 3.081192168 | 0.025685788 | |
| St8sia6 | 2.556608959 | 0.000223695 | |
| B430306N03Rik | 2.48050793 | 0.035173654 | Yes |
| Id3 | 2.465815673 | 0.019139609 | |
| Pik3ip1 | 2.239096221 | 0.02553885 | |
| Gm11808 | 2.201610226 | 0.001790636 | |
| Pde3b | 2.116019659 | 0.022864302 | |
| St8sia1 | 2.101573053 | 0.002901057 | Yes |
| Rpl39 | 1.987335644 | 0.018638235 | |
| Rps27 | 1.98443294 | 0.003529233 | |
| Rpl23a | 1.956213335 | 0.002832813 | |
| Gm10073 | 1.953001613 | 0.005186147 | |
| Il7r | 1.948327528 | 0.000159169 | Yes |
| Rps24 | 1.947715696 | 1.54E-05 | |
| Tomm7 | 1.945285451 | 0.041101661 | |
| Gm13212 | 1.926314285 | 0.046383967 | |
| Rps19 | 1.870594769 | 8.55E-05 | Yes |
| Rpl37 | 1.868758872 | 0.000884253 | |
| Rpl9-ps6 | 1.86387934 | 0.005186147 | |
| Ccr7 | 1.852664747 | 0.039075329 | |
| Rps23 | 1.829116323 | 0.00148382 | |
| Rps29 | 1.828263829 | 0.004129927 | |
| Rps7 | 1.819247973 | 0.000884253 | |
| Rpl35 | 1.815845372 | 0.004849943 | |
| Rps20 | 1.796811133 | 0.003529233 | |
| Rps21 | 1.791878082 | 0.022274648 | |
| Lef1 | 1.782536087 | 0.015427684 | |
| Rpl36a | 1.781657838 | 0.006552298 | |
| Bcl2l11 | 1.779317672 | 0.019139609 | Yes |
| Itm2a | 1.771539743 | 0.035173654 | |
| Sit1 | 1.768058047 | 0.022864302 | |
| Rpl37a | 1.763991694 | 0.029314942 | |
| Rpl30 | 1.739178854 | 0.001424461 | |
| Myb | 1.734483944 | 0.04984297 | |
| Rps10 | 1.731510725 | 0.002831294 | |
| Rps14 | 1.723300596 | 0.007862519 | |

| | | | |
|---|---|---|---|
| Cd1d1 | 1.723060393 | 0.035712477 | Yes |
| Rps15a | 1.711325013 | 0.00026822 | Yes |
| Rpl35a | 1.707402838 | 0.019240761 | |
| Rps27a | 1.700188841 | 0.00034891 | |
| Fau | 1.688269829 | 0.002536583 | |
| Rpl34 | 1.677755734 | 0.001989184 | |
| Arl5c | 1.673217245 | 0.036633277 | |
| Dnaja1 | 1.669952781 | 0.040200656 | |
| Rps17 | 1.667966886 | 0.004068099 | |
| Rps12 | 1.663058828 | 0.003529233 | |
| Rps13 | 1.660779053 | 0.005944903 | |
| Rpl27a | 1.649809416 | 0.008859978 | |
| Tmem55b | 1.648571096 | 0.0373 | Yes |
| Fkbp5 | 1.647501334 | 0.007563338 | |
| Rps4x | 1.635591834 | 0.000263015 | |
| Ppp1r21 | 1.627007854 | 0.018371985 | |
| Rps16 | 1.621100227 | 0.00148382 | |
| Rpl38 | 1.613333243 | 0.019240761 | |
| Rpl27 | 1.606096467 | 0.038418338 | |
| Rpl31 | 1.59197943 | 0.024387894 | |
| Rps25 | 1.570858837 | 0.007862519 | |
| Rpl23 | 1.568420293 | 0.004129927 | |
| Rps18 | 1.564795739 | 0.008788791 | |
| Rps5 | 1.547865453 | 0.013312125 | |
| Rpl32 | 1.542070794 | 0.003529233 | |
| Rps3a1 | 1.541151816 | 0.004443923 | |
| Npc2 | 1.534808143 | 0.025685788 | |
| Foxp1 | 1.529301483 | 0.040200656 | Yes |
| Rps9 | 1.526465121 | 1.54E-05 | |
| Rpl17 | 1.520313901 | 0.00148382 | |
| Pdcd4 | 1.506918002 | 0.006210999 | |
| Nacc1 | 0.665779419 | 0.022864302 | |
| Top2a | 0.664600916 | 0.029314942 | |
| Mcm5 | 0.650790829 | 0.039075329 | |
| Slc25a24 | 0.642007235 | 0.023530592 | |
| Hivep3 | 0.634441268 | 0.019139609 | |
| Nucb1 | 0.628729545 | 0.03599535 | |
| Lmf2 | 0.628260221 | 0.038233226 | |
| Uhrf1 | 0.62055072 | 0.019390571 | |
| Anxa2 | 0.608148914 | 0.029314942 | |
| Cxcr3 | 0.603697435 | 0.000101658 | |
| Fbxw8 | 0.591397429 | 0.035794654 | |
| Cacna1d | 0.563353747 | 0.036633277 | |

| | | |
|---|---|---|
| F2r | 0.549627748 | 0.003341628 |
| Ybx3 | 0.547301238 | 0.036633277 |
| Scd2 | 0.540536083 | 0.000884253 |
| Twsg1 | 0.539781658 | 0.041440095 |
| Kif5c | 0.523072074 | 0.011318627 |
| Ccl5 | 0.520537077 | 0.00148382 |
| Il21 | 0.513454947 | 0.029314942 |
| Cdh23 | 0.508131895 | 0.04315034 |
| Itga7 | 0.48519738 | 0.038233226 |
| Chst2 | 0.466386912 | 0.01081644 |
| Spag5 | 0.464563332 | 0.036633277 |
| Cpne7 | 0.454809801 | 0.008159095 |
| Susd2 | 0.45381711 | 0.023284384 |
| Nkg7 | 0.431360675 | 0.004644651 |
| Angptl2 | 0.417742475 | 9.68E-05 |
| Klc3 | 0.405319194 | 0.040200656 |
| Vdr | 0.399663505 | 0.026146195 |
| Spp1 | 0.374053736 | 0.001989184 |
| Fbn2 | 0.367838128 | 0.002831294 |
| Gzmb | 0.366405359 | 0.029314942 |
| Camk2b | 0.357406479 | 0.002536583 |
| Pdgfb | 0.344893546 | 0.027105042 |
| Myh3 | 0.342846996 | 0.044192138 |
| Ecel1 | 0.321449636 | 0.008159095 |
| Chn1 | 0.199631249 | 0.003529233 |
| Zfp111 | 0.175301653 | 0.042117349 |
| Cdkn3 | 0.151773154 | 0.023284384 |
| Col5a3 | 0.072305597 | 0.000884253 |
| Nme2 | 0.023094432 | 0.038233226 |
| Lrtm2 | 0.019399107 | 0.047165458 |
| Wdr93 | 0.017296657 | 0.028994523 |
| Col4a2 | 0.016373643 | 0.035173654 |
| Ulk4 | 0.015490496 | 0.016120476 |
| Prg2 | 0.015404828 | 0.023284384 |

**Table S4. Significantly altered pathways based on gene expression changes in splenic CD4⁺ T cells *in vivo* treated with LNA-22.**

| # | Maps | In Data | Total | p-value | Network Objects from Active Data |
|---|---|---|---|---|---|
| 1 | Populations of skin dendritic cells involved in contact hypersensitivity | 3 | 18 | 7.13E-05 | CD36, CCR7, CD1d |
| 2 | Cell cycle_Transition and termination of DNA replication | 3 | 25 | 1.96E-04 | TOP2 alpha, Ubiquitin, TOP2 |
| 3 | Proteolysis_Putative SUMO-1 pathway | 3 | 29 | 3.08E-04 | c-Myb, Ubiquitin, TOP2 |
| 4 | Memory CD8+ T cells in allergic contact dermatitis | 3 | 40 | 8.04E-04 | CCL5, IL7RA, CCR7 |
| 5 | Role of platelets in the initiation of in-stent restenosis | 3 | 43 | 9.94E-04 | CCL5, PAR1, PDGF-B |
| 6 | Immune response_Role of DPP4 (CD26) in immune regulation | 3 | 47 | 1.29E-03 | CCL5, Ubiquitin, Granzyme B |
| 7 | Immune response_T cell subsets: cell surface markers | 3 | 52 | 1.73E-03 | IL7RA, CXCR3, CCR7 |
| 8 | Neurophysiological process_Dynein-dynactin motor complex in axonal transport in neurons | 3 | 54 | 1.93E-03 | Ubiquitin, Kinesin light chain, Kinesin heavy chain |
| 9 | Transcription_Role of VDR in regulation of genes involved in osteoporosis | 3 | 61 | 2.74E-03 | VDR, CYP27B1, Osteopontin |
| 10 | Signal transduction_mTORC2 upstream signaling | 3 | 65 | 3.28E-03 | RPL23a, RPS16, RPL23 |
| 11 | Schema: Initiation of T cell recruitment in allergic contact dermatitis | 2 | 19 | 3.37E-03 | CXCR3, CD1d |
| 12 | Development_WNT signaling pathway. Part 1. Degradation of beta-catenin in the absence WNT signaling | 2 | 19 | 3.37E-03 | Ubiquitin, Tcf(Lef) |
| 13 | Eosinophil adhesion and transendothelial migration in asthma | 3 | 68 | 3.73E-03 | CCL5, Osteopontin, Collagen IV |
| 14 | Signal transduction_mTORC2 downstream signaling | 3 | 68 | 3.73E-03 | IL7RA, Bim, FBX29 |
| 15 | Mechanisms of drug resistance in SCLC | 3 | 70 | 4.04E-03 | TOP2 alpha, Osteopontin, Collagen IV |
| 16 | Blood coagulation_Platelet microparticle generation | 3 | 72 | 4.38E-03 | CCL5, MyHC, PAR1 |

| | | | | | |
|---|---|---|---|---|---|
| 17 | Mast cell migration in asthma | 3 | 73 | 4.55E-03 | CCL5, CXCR3, CCR7 |
| 18 | Action of GSK3 beta in bipolar disorder | 2 | 23 | 4.93E-03 | Tcf(Lef), Lef-1 |
| 19 | G protein-coupled receptors signaling in lung cancer | 3 | 76 | 5.09E-03 | CCL5, Galpha(i)-specific peptide GPCRs, Galpha(q)-specific peptide GPCRs |
| 20 | Immune response_T cell subsets: secreted signals | 2 | 25 | 5.81E-03 | CCL5, IL-21 |
| 21 | HSP70 and HSP40-dependent folding in Huntington's disease | 2 | 25 | 5.81E-03 | Ubiquitin, Hdj-2 |
| 22 | Defective macrophage-mediated bacterial phagocytosis in COPD | 2 | 25 | 5.81E-03 | CD36, SR-BI |
| 23 | Inhibition of TGF-beta signaling in gastric cancer | 2 | 25 | 5.81E-03 | Bim, Ubiquitin |
| 24 | Pioglitazone and Rosiglitazone in treatment of type 2 diabetes and metabolic syndrome X | 2 | 26 | 6.28E-03 | CD36, SR-BI |
| 25 | Role of fibroblasts and keratinocytes in the elicitation phase of allergic contact dermatitis | 2 | 26 | 6.28E-03 | CCL5, CXCR3 |
| 26 | Th1 and Th17 cells in an autoimmune mechanism of emphysema formation in smokers | 2 | 28 | 7.26E-03 | CXCR3, Osteopontin |
| 27 | Impaired inhibition of Th17 cell differentiation by IFN-beta in multiple sclerosis | 2 | 28 | 7.26E-03 | IL-21, Osteopontin |
| 28 | Inter-cellular relations in COPD (general schema) | 2 | 30 | 8.31E-03 | CXCR3, Granzyme B |
| 29 | Glomerular injury in Lupus Nephritis | 3 | 92 | 8.64E-03 | Annexin II, CCL5, PDGF-B |
| 30 | Role of IL-2 in the enhancement of NK cell cytotoxicity in multiple sclerosis | 2 | 31 | 8.86E-03 | Ubiquitin, Granzyme B |
| 31 | Deregulation of canonical WNT signaling in major depressive disorder | 2 | 31 | 8.86E-03 | Tcf(Lef), Lef-1 |
| 32 | Stem cells_Cooperation between Hedgehog, IGF-2 and HGF signaling pathways in medulloblastoma stem cells | 2 | 32 | 9.42E-03 | Tcf(Lef), Lef-1 |
| 33 | The role of KEAP1/NRF2 pathway in skin sensitization | 2 | 32 | 9.42E-03 | Ubiquitin, CCR7 |
| 34 | Interleukins-induced inflammatory signaling in normal and asthmatic airway epithelium | 2 | 32 | 9.42E-03 | CCL5, CXCR3 |

| | | | | | |
|---|---|---|---|---|---|
| 35 | Stem cells_Role of PKR1 and ILK in cardiac progenitor cells | 2 | 33 | 1.00E-02 | Tcf(Lef), Lef-1 |
| 36 | NK cells in allergic contact dermatitis | 2 | 33 | 1.00E-02 | CCL5, CXCR3 |
| 37 | Extracellular matrix-regulated proliferation of airway smooth muscle cells in asthma | 2 | 35 | 1.12E-02 | PDGF-B, Collagen IV |
| 38 | Mechanisms of CAM-DR in multiple myeloma | 2 | 35 | 1.12E-02 | Bim, TOP2 alpha |
| 39 | Langerhans cell migration to lymph nodes in allergic contact dermatitis | 2 | 35 | 1.12E-02 | CCR7, Collagen IV |
| 40 | Development_Hedgehog and PTH signaling pathways in bone and cartilage development | 2 | 36 | 1.18E-02 | VDR, Osteopontin |
| 41 | WNT signaling in gastric cancer | 2 | 36 | 1.18E-02 | Ubiquitin, Tcf(Lef) |
| 42 | Th17 cytokines in COPD | 2 | 36 | 1.18E-02 | CXCR3, IL-21 |
| 43 | iNKT cell-keratinocyte interactions in allergic contact dermatitis | 2 | 36 | 1.18E-02 | CXCR3, CD1d |
| 44 | Immune response_Generation of memory CD4+ T cells | 2 | 37 | 1.25E-02 | IL7RA, CCR7 |
| 45 | Cell adhesion_Tight junctions | 2 | 37 | 1.25E-02 | Tcf(Lef), CSDA |
| 46 | Eosinophil-derived cytokines in airway remodeling in asthma | 2 | 38 | 1.31E-02 | PDGF-B, Osteopontin |
| 47 | Regulation of degradation of deltaF508-CFTR in CF | 2 | 39 | 1.38E-02 | Ubiquitin, Hdj-2 |
| 48 | VLDL, LDL dyslipidemia in type 2 diabetes and metabolic syndrome X | 2 | 39 | 1.38E-02 | CD36, SR-BI |
| 49 | PXR-mediated direct regulation of xenobiotic metabolizing enzymes / Rodent version | 2 | 39 | 1.38E-02 | CD36, SR-BI |
| 50 | Role of iNKT and B cells in T cell recruitment in allergic contact dermatitis | 2 | 39 | 1.38E-02 | CXCR3, CD1d |
